# Co-Essentiality Analysis Identifies PRR12 as a Regulator of Cohesin and Genome Integrity

**DOI:** 10.1101/2024.03.29.587394

**Authors:** Alexandra L. Nguyen, Eric Smith, Iain M. Cheeseman

## Abstract

The cohesin complex is critical for genome regulation, relying on specialized co-factors to mediate its diverse functional activities. Here, by analyzing patterns of similar gene requirements across cell lines, we identify PRR12 as a regulator of cohesin and genome integrity. We show that PRR12 interacts with cohesin and PRR12 loss results in a reduction of nuclear-localized cohesin and an accumulation of DNA lesions. We find that different cell lines across human and mouse exhibit significant variation in their sensitivity to PRR12 loss. Unlike the modest phenotypes observed in human cell lines, PRR12 depletion in mouse cells results in substantial genome instability. Despite a modest requirement in human cell lines, mutations in PRR12 lead to severe developmental defects in human patients, suggesting context-specific roles in cohesin regulation. By harnessing comparative studies across species and cell lines, our work reveals critical insights into how cohesin is regulated across diverse cellular contexts.

## Introduction

The precise organization and regulation of the genome is critical for cellular function. In human cells, over 6 feet of linear DNA must be compressed into a nucleus of only ∼10 μm in diameter. Despite the need to organize and compact the genome to only a fraction of its length, the genome must remain dynamic and accessible to allow for key regulatory processes including gene expression, DNA repair, DNA replication, and cell division. The ring-like cohesin complex performs multiple essential functions in the cell that contribute to overall genome organization and function [1–3]. Cohesin is comprised of two structural maintenance of chromosome proteins (SMC1, SMC3) and a Kleisin subunit (Rad21) [4–6]. In addition to the core cohesin complex proteins, a STAG protein (STAG1, STAG2, or STAG3) interacts with the complex and is required for the association of cohesin with the DNA [7–9]. In mitosis, cohesin acts to ensure that sister chromatids remain physically connected until anaphase onset, a function known as sister chromatid cohesion [10, 11]. However, throughout the cell cycle, cohesin contributes to diverse aspects of genome structure, organizing DNA into loops that are important for both DNA organization and gene expression [12–16]. Finally, cohesin contributes to DNA replication [17–21], and post-replicative DNA repair [22–25]. The diverse functional activities of the cohesin complex depend on its association with multiple co-factor proteins that regulate cohesin localization, stability, and function in a context-specific manner (Fig. 1) [2, 26, 27]. For example, cohesin is stabilized on DNA via its association with NIPBL and MAU2 [27–30]. Recent work also identified the NIPBL/MAU2 complex as a critical regulator of the cohesin loop extrusion process [31]. Additional regulators further contribute to cohesin establishment, maintenance, stability and removal (Fig. 1) [2, 26, 32].

**Figure 1.**
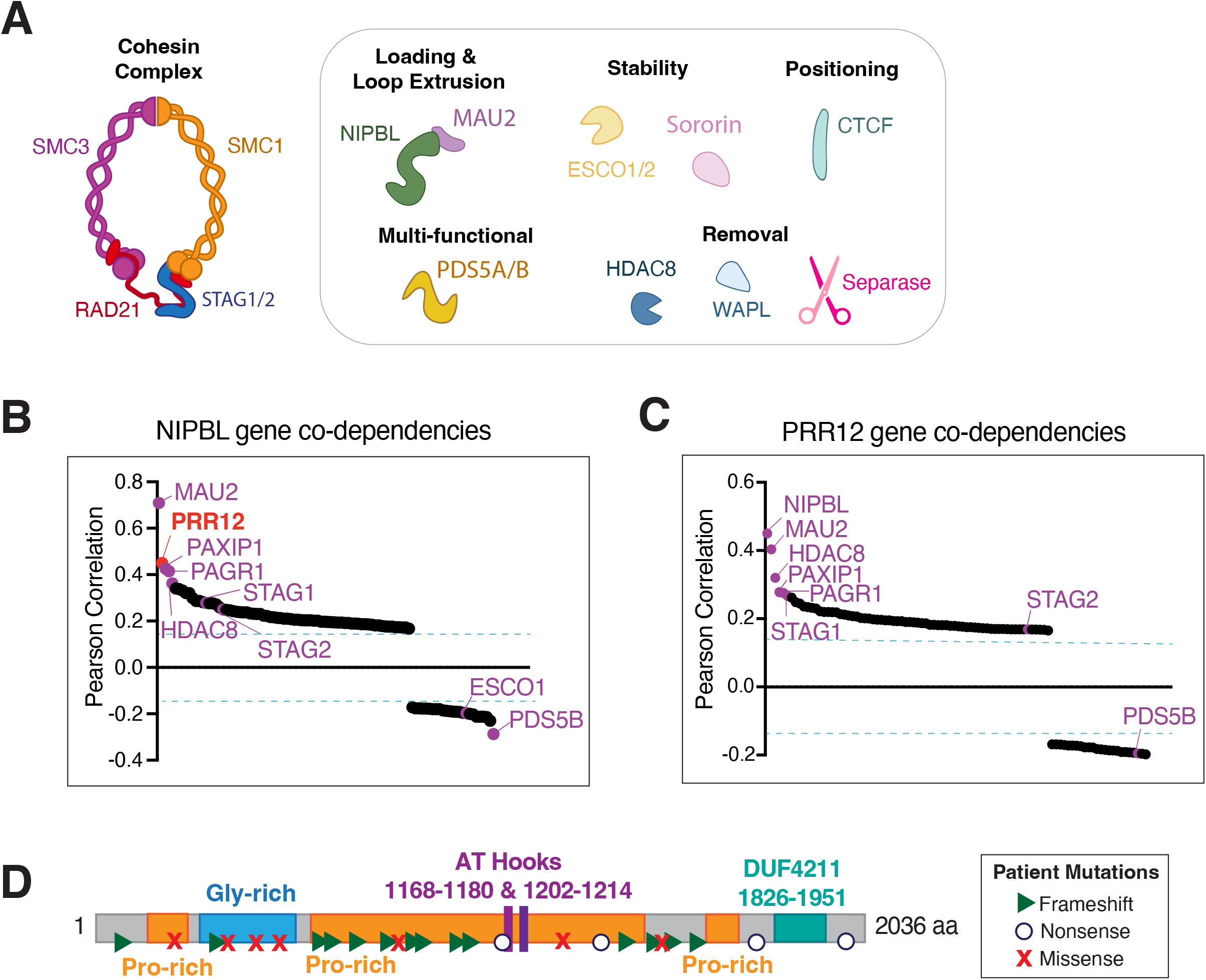
Identification of PRR12 through co-essentiality with multiple cohesin regulatory proteins. (A) Diagram of the cohesin complex and associated co-factor proteins. (B-C) Gene co-dependency predictions from DepMap. Scores represent the Pearson correlation coefficients of CRISRP gene effects scores between each gene pair. Previously-identified cohesin regulatory proteins are indicated in purple. B) Gene co-dependency predictions for NIPBL. C) Gene co-dependency predictions for PRR12. (D) Graphic depicting the protein structure for PRR12 and disease-associated mutations identified in human patients. Symbols represent the mutation type (Nonsense, Missense, Frameshift).

The complete loss-of-function of the cohesin complex results in cellular and organismal lethality [33]. However, mutations in the genes that regulate cohesin can result in developmental and neurological disorders in human patients, diseases collectively known as cohesinopathies [33–35]. Cornelia de Lange syndrome (CdLS), the most well-known cohesinopathy, results from mutations in NIPBL and is characterized by severe cognitive delays, craniofacial and limb dysmorphism, cardiac defects, and growth delays [33, 34, 36–39]. Similarly, Roberts syndrome, which exhibits comparable developmental and cognitive delay phenotypes as CdLS, results from mutations in the cohesin acetyltransferase ESCO2 [40, 41]. Despite the critical importance for cohesin regulatory factors for human development and health, loss of many of these proteins in standard human cell culture systems often results in only mild phenotypic consequences [42]. One explanation for these discrepancies may be the situation-specific roles that the cohesin complex plays in genome regulation that are not fully captured using standard cell lines grown in culture. These challenges have posed significant limitations to our understanding of how cohesin is regulated and the precise mechanisms by which its co-factors function. Additionally, the limited number of co-factors identified to date cannot account for all of the known functions of cohesin, suggesting additional regulators remain to be discovered. Differences in cellular origin, gene expression programs, or a cell’s mutational spectrum can create varying requirements and sensitivities for proteins across distinct cell lines. By exploiting these differences, it is possible to identify specific situations in which cohesin co-factor proteins become essential for cellular viability, and utilize these situations to analyze their function.

Here, we harnessed the differential gene requirements of cohesin co-factor proteins in combination with gene co-essentiality analysis to identify genes that function in cohesin complex regulation. By comparing differential genetic dependencies across diverse cell lines with established cohesin regulators, we identified the functionally-uncharacterized protein PRR12 as a regulator of cohesin and genome integrity. We find that PRR12 interacts with the cohesin complex, and that the loss of PRR12 results in reduced nuclear-localized cohesin and a substantial increase in DNA double-strand breaks. Importantly, we show that the requirement for PRR12 differs significantly across cell lines, with human HeLa cells showing reduced sensitivity to PRR12 loss in comparison to mouse NIH-3T3 cells, which display dramatic defects in genome integrity in the absence of PRR12. Together, these data highlight the power of utilizing a comparative genomics approach for the identification and investigation of cohesin regulatory mechanisms.

## Results

### Identification of PRR12 through co-essentiality with multiple cohesin regulatory proteins

Recent large-scale analyses using systematic CRISPR/Cas9-based functional genetics profiling has analyzed gene requirements across more than 1000 different human cell lines [42]. In addition to revealing the functional requirements for a specific gene target in individual cell lines, this analysis reveals patterns of gene sensitivities, where the degree to which a gene is essential for proliferation or survival varies in a cell line-dependent manner. These patterns of genetic requirements can be used predict gene function, as genes that function in shared molecular pathways are predicated to display similar requirements [43]. Here, we harnessed these co-essentiality predictions as a means to identify previously uncharacterized genes that participate in core cellular processes. We focused our analysis on the cohesin pathway, which participates in genome organization, gene expression, cell division, and genomic integrity. Using CRISPR scores from Project Achilles (DepMap), in which a negative score reflects a requirement for a gene target for cellular fitness, we analyzed correlations in gene requirements across human cell lines [42]. In this manner, we used established cohesin factors as “baits” to identify previously unknown co-functional cohesin regulators. For example, NIPBL is an established cohesin regulator that participates in DNA loop extrusion and DNA entrapment [29, 31, 44–46]. Using NIPBL as a bait, we observed multiple established cohesin regulators as displaying strongly correlated gene requirements across cell lines including the NIPBL-binding partner, MAU2, which displays a correlation of 0.70 (Fig. 1A-B). Interestingly, in addition to established players in cohesin biology, we identified the previously uncharacterized gene, PRR12, as the second strongest hit, with a correlation score of 0.44. In a reciprocal analysis, co-essentiality correlations for PRR12 revealed strong relationships with multiple known cohesin regulators (Fig. 1A,C), providing support for a role for PRR12 in a cohesin-related pathways.

PRR12 is a large 2036 residue protein that is comprised of proline and glycine-rich regions, as well as two predicted AT hook motifs and an annotated domain of unknown function (DUF) (Fig. 1D). The primary amino acid sequence for PRR12 is highly conserved across vertebrate organisms, including 93.6% amino acid identity between the mouse and human PRR12 proteins. Although the cellular functions of PRR12 have not be analyzed previously, mutations in PRR12 lead to severe developmental and neurological disorders in humans [47–50] (Fig. 1D). Notably, the resulting phenotypes observed in human patients with PRR12 mutations are highly similar to those reported when known cohesin regulators are perturbed, for example in Cornelia de Lange Syndrome which results from mutations in NIPBL [34, 47, 50–52]. Thus, based on co-varying functional requirements with established players in cohesin biology and these reported patient mutations, we chose to investigate the cellular function of PRR12.

### PRR12 co-localizes with the cohesin complex

To investigate the relationship between PRR12 and cohesin, we first sought to analyze the localization of PRR12 in cells. The nuclear-localized cohesin complex displays specific spatial and temporal dynamics that are critical to its function, with lower levels in G1 when cohesin is most dynamic, increased levels in S phase and G2 as cohesin becomes entrapped onto DNA to establish cohesive cohesin, and then reduced levels in mitosis due to the removal of the majority of cohesin through the WAPL-based prophase pathway [53–57]. Similar to the observed localization of cohesin [58], ectopically-expressed GFP-PRR12 localized to the nucleus in interphase cells, is undetectable on DNA in mitosis, and rapidly re-localized to the nucleus as daughter cells enter G1 (Fig. 2A).

**Figure 2.**
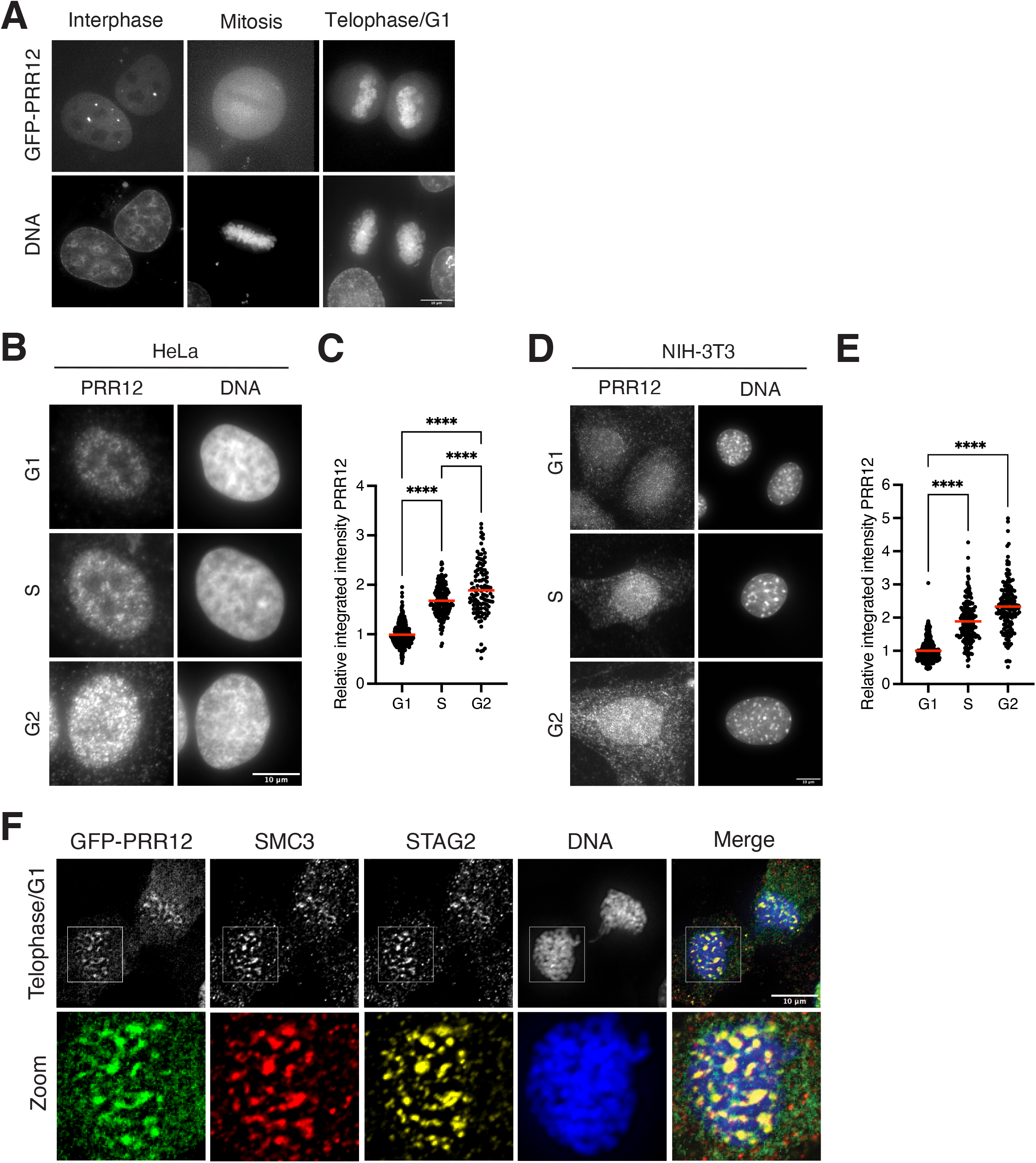
PRR12 co-localizes with the cohesin complex. (A) Representative Z-projected images of HeLa cells ectopically expressing human GFP-PRR12 at the indicated cell stages. Images show GFP-PRR12 and DNA (Hoechst). (B) Representative Z-projected images of HeLa cells synchronized at the indicated cell stages. Images show anti-PRR12 antibody and DNA (Hoechst). (C) Relative pixel intensity for nuclear localized PRR12, quantified from B. Values are normalized to the average control value within each experiment. *n* = approximately 150 cells per condition, across 2 experimental replicates. (D) Representative Z-projected images of NIH-3T3 cells synchronized at the indicated cell stages. Images show anti-PRR12 antibody and DNA (Hoechst). (E) Relative pixel intensity for nuclear localized PRR12, quantified from D. *n* = approximately 150 cells per condition, across 2 experimental replicates. (F) Representative Z-projected images of late telophase/G1 stage NIH-3T3 cells ectopically expressing human GFP-PRR12. Images show GFP-PRR12 (green), anti-SMC3 (yellow), anti-STAG2 (red), and DNA (Hoechst). Boxes indicate zoom. Red bars indicate mean. Error bars indicate SD. One-way ANOVA was performed (**** = <0.0001). Scale bars, 10 μM.

To further assess the cell cycle dynamics of PRR12 localization, we utilized an antibody against PRR12 and a cell synchronization approach to quantify the nuclear localization of PRR12 in G1, S, and G2. Similar to cohesin, we found that the levels of nuclear-localized PRR12 changed across the cell cycle in Hela cells, with a basal level in G1 followed by a significant nuclear accumulation of PRR12 in S and G2 (Fig. 2B-C). Based on immunofluorescence with the anti-PRR12 antibody, we also observed a similar localization behavior and cell cycle dynamics in mouse NIH-3T3 cells (Fig. 2D-E).

The localization of PRR12 to the nucleus is consistent with a role for PRR12 in genome function. However, as cohesin and PRR12 are both nuclear-localized in interphase similar to many DNA-associated proteins, it is challenging to detect precise co-localization. We therefore analyzed the localization of the cohesin complex and PRR12 in late telophase/G1 cells, a timepoint during which both cohesin and PRR12 can be detected re-localizing to the DNA in discrete populations (Fig. 2F). By analyzing the localization of GFP-PRR12 and the core cohesin components, SMC3 and STAG2, in late telophase/G1 NIH-3T3 cells, we observed close co-localization of all three proteins (Fig. 2F). Together, this co-localization analysis is consistent with a relationship between PRR12 and the cohesin complex.

### PRR12 interacts with cohesin and its co-factors via its C-terminus

The co-localization of PRR12 with cohesin throughout the cell cycle suggests that these proteins may interact. To test this, we conducted affinity purifications coupled with mass spectrometry to identify the proteins that interact with PRR12 in cells. To this end, we isolated PRR12 via GFP immunoprecipitation from both human Hela and mouse NIH-3T3 cell lines stably expressing human GFP-PRR12. In both human and mouse cells, GFP-PRR12 immunoprecipitated with all established subunits of the cohesin complex in addition to multiple cohesin-associated proteins, including NIPBL and MAU2 (Fig. 3A). Notably, multiple known cohesin interacting proteins, such as components of the MCM Replication complex and proteins in the DNA damage response pathways, were also identified in both samples. Consistent with this interaction, prior work also isolated PRR12 in large-scale cohesin immunoprecipitation experiments [59].

**Figure 3.**
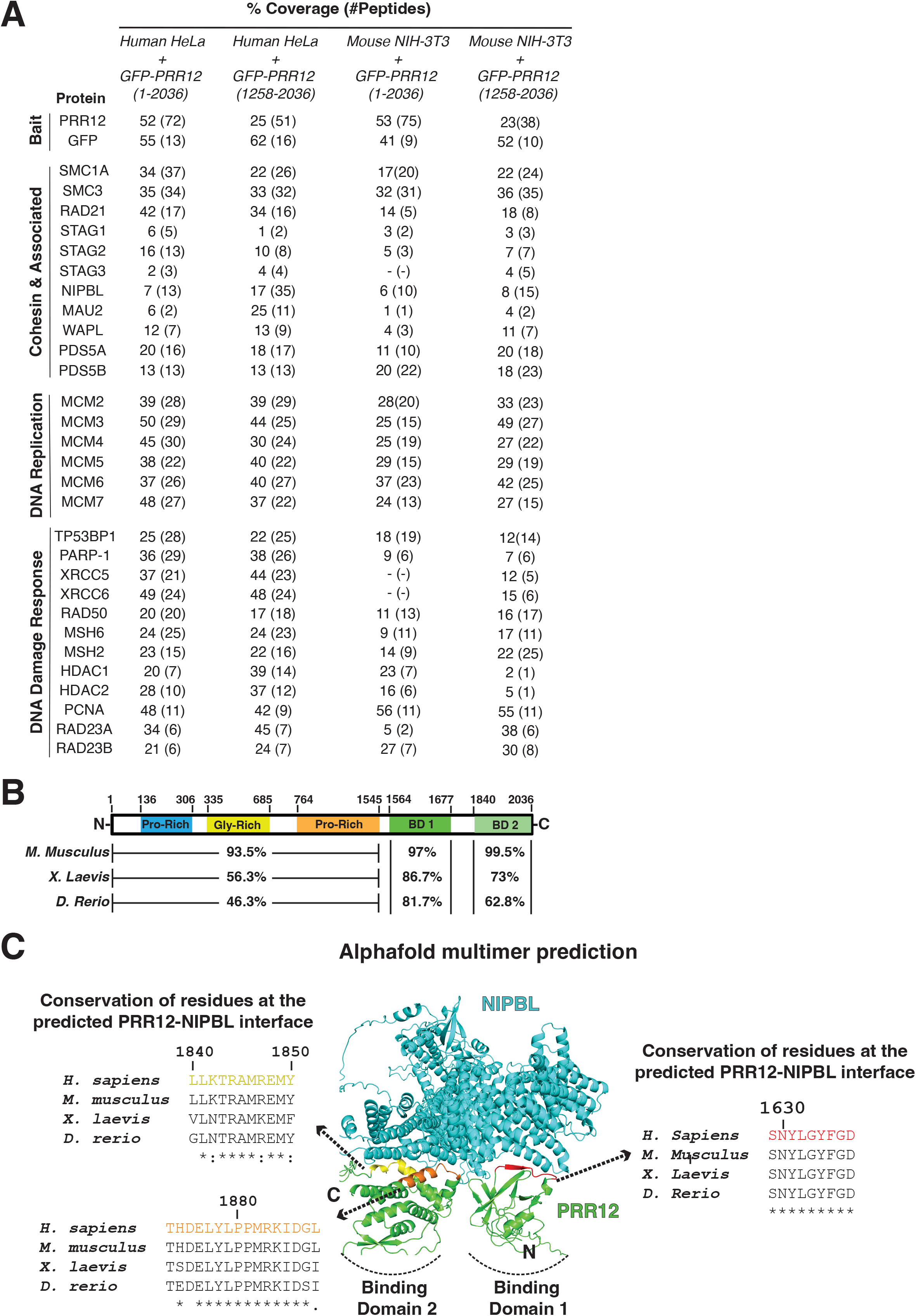
PRR12 interacts with cohesin and its co-factors via its C terminus. (A) Mass spectrometric analysis from immunoprecipitation experiments using the indicated fusion proteins (italicized) in human HeLa and mouse NIH-3T3 cell lines. Percent sequence coverage and the number of peptides obtained for each protein identified is shown. (B) Diagram of PRR12 protein domain structure with indicated percent identity across species. Predicted interactions domains are indicated. (C) Alphafold multimer predictions for PRR12 in complex with NIPBL. Protein residue conservation in the predicted binding sites are indicated. See also supplemental Figure 1.

To understand which domains in PRR12 are involved in the interaction with the cohesin complex, we conducted structural predictions using Alphafold [60]. The N-terminal 1564 residues of PRR12 are predicted to be largely unstructured, with the potential presence of polyproline helices due to the large number of proline residues. The C-terminus of PRR12 is predicted to have two highly-conserved, structured domains that we will refer to as binding domain 1 (residues 1564-1677) and binding domain 2 (residues 1840-2036) (Fig. 3B and Supplemental Fig. 1A-C). Binding domain 1 is predicted to be composed of 8 contorted anti-parallel beta-sheets and two helices whereas binding domain 2 is predicted to contain a bundle of 7 helices flanked by 3 anti-parallel beta-sheets (Supplemental Fig. 1A). Given the strong co-essentiality of PRR12 with NIPBL and the identified interaction between NIPBL-PRR12 in our affinity purifications, we used Alphafold-Multimer to predict the structure of a potential PRR12-NIPBL complex (Fig. 3C)[61]. We found that two highly conserved regions in PRR12, one in binding domain 1 (residues 1629-1637) and the other in binding domain 2 (residues 1840-1850 and 1873-1888), are predicted to be located at the interface of PRR12 and NIPBL (Fig. 3C).

To test if the C-terminal region of PRR12 is sufficient for the direct interaction between PRR12 and NIPBL or the cohesin complex, we stably expressed a GFP-tagged construct of the C-terminus of human PRR12 in both human HeLa and mouse NIH-3T3 cells and performed immunoprecipitation experiments as described above. These immunoprecipitations revealed that C-terminal region of PRR12 (residues 1258-2036; GFP-C-term. PRR12) was sufficient to mediate its associations with NIPBL, cohesin, and the other associated co-factors (Fig. 3A), suggesting that this constitutes its core interacting domain. Overall, these results demonstrate that PRR12 interacts with the cohesin complex in cells and suggest this interaction may be facilitated in part by the NIPBL/MAU2 complex.

### PRR12 is required for viability in a cell type-specific manner

Analysis of varying gene requirements across different human cell lines indicated that PRR12 displays a similar pattern of dependencies as multiple cohesin regulatory proteins (Fig. 1B-C) [42]. In these analyses, the functional requirement or “essentiality” for a given gene is calculated based on the relative change in proliferation under competitive growth conditions. CRISPR gene scores less than −0.5 represent fitness-conferring genes in most cell lines, whereas scores less than −1 are consistent with significant lethality. We found that the requirement for PRR12 varied across human cell lines. However, in each case, the corresponding effect on cellular fitness following the loss of PRR12, as reflected by the CRISPR score, suggests that PRR12 is largely dispensable for viability in human cell lines (Fig. 4A). In contrast, we found that large-scale CRISPR/Cas9-based functional screens in mouse cells revealed a highly sensitized requirement for PRR12 (Fig. 4B). We observed a similar pattern of behavior for multiple cohesin co-factors, with more modest fitness requirements in human cells, but substantially increased requirements in mouse cells (Fig. 4A-B).

**Figure 4.**
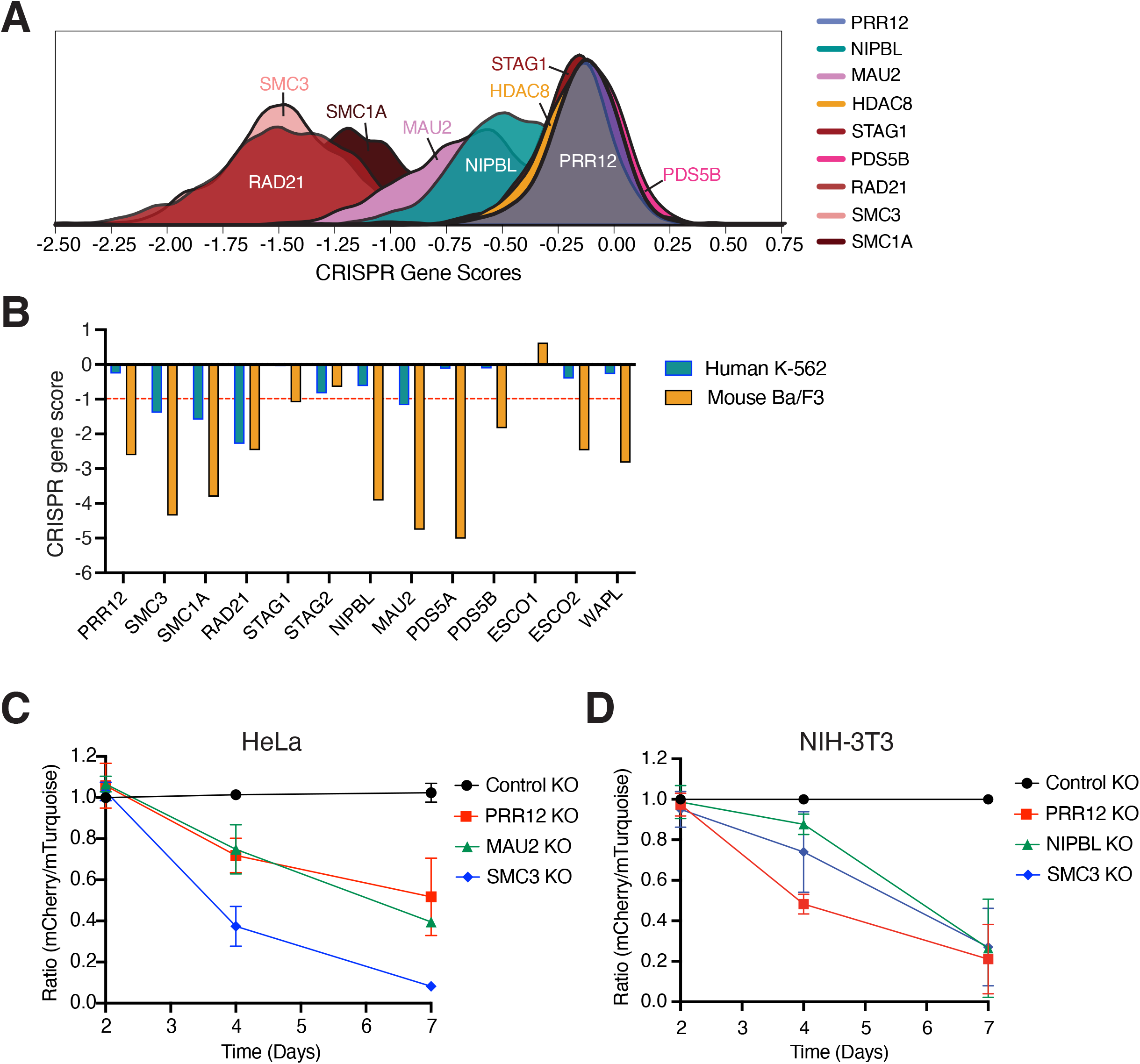
PRR12 is required for viability in a cell type-specific manner. (A) Diagram depicting CRISPR gene effect scores from DepMap for PRR12 and known cohesin regulators across human cell lines. (B) Comparison of CRISPR gene effect scores for PRR12 and known cohesin regulators in human K-562 (DepMap) and mouse BaF3 cell lines [83]. (C-D) Competitive proliferation assay in human Hela (C) and mouse NIH-3T3 cell lines (D) transduced with lentivirus containing the indicated mCherry-expressing gene knockout plasmids and a control mTurquoise-expressing sgAAVS1 control gene knockout plasmid. At day 2 after virus transduction, mCherry and mTurquoise -expressing cells were mixed 1:1 and proliferation was monitored by flow cytometry analysis. The ratio of mCherry to mTurquoise-positive cells at the indicated timepoint was quantified and normalized to the control mCherry-expressing sgAAVS1 control vs mTurquoise-expressing sgAAVS1 control combined group at each time point. Error bars represent the SD for growth rates across 3 experimental replicates.

To test these behaviors and the differing requirements for PRR12 and cohesin regulatory proteins between human and mouse cells directly, we used a cellular competition assay to test the relative growth of control cells and knockout cells that are co-cultured for a direct comparison for their fitness behavior. In human HeLa cells, SMC3 knockout cells were rapidly lost from the population, with 63% of knockout cells lost from the population by day 4 of gene knockout and 99% of knockout cells lost by day 7 (Fig. 4C). These data are consistent with the documented essential role for SMC3 in cellular viability. In contrast, PRR12 and MAU2 knockout cells displayed a more modest fitness defect. We observed a reduced fraction of PRR12 or MAU2 knockouts relative to controls, with 48% and 61% fewer cells respectively, observed at the end of the 7-day time course. In contrast, using a similar cellular competition assay in mouse NIH-3T3 cells, we found that mouse cells were highly sensitive to the loss of the cohesin subunit RAD21, NIBPL, and PRR12 with similar growth dynamics throughout the time course. By day 7 of the time course, 79%, 74% and 73% of PRR12, NIPBL, and RAD21 knockout cells had been lost, respectively (Fig. 4D).

The strong requirement for PRR12 In mouse cells provides a situation under which PRR12 is specifically required for cell growth and viability. As the PRR12 protein is highly conserved between mouse and humans (Fig. 3B) and displays similar localization behavior and protein interactions between mouse and human cell lines (Fig. 2B-E), this provides a powerful strategy to assess PRR12 function and its potential role in cohesin biology. To define the functional contributions of PRR12 and provide insights into why this protein is differentially required across cell lines, we therefore choose to conduct comparative functional studies in both mouse NIH-3T3 cells and human HeLa cells.

### PRR12 is required for cohesin localization

Using a CRISPR/Cas9-based conditional gene knockout approach, we next sought to define the consequences of PRR12 loss. Analysis of PRR12 knockout NIH-3T3 cells revealed severe defects in DNA organization and genome integrity including the presence of micronuclei (32.5%) and bi-nucleated cells (9.5%) (Fig. 5A-B). Consistent with the predicted co-dependency between PRR12 and cohesin co-factors, knockout of NIPBL (22.2% micronuclei, 5.8% binucleation) and RAD21 (31.3% micronuclei, 10.9% binucleation) also resulted in similar DNA morphology defects (Fig. 5A-B). Knockout of PRR12 in HeLa cells also resulted in DNA abnormalities, but these phenotypes were significantly reduced in their penetrance and severity relative to PRR12 knockout NIH-3T3 cells (Fig. 5C). Notably, knockout of MAU2 resulted in a similar percentage of micronuclei and binucleated cells as knockout of PRR12 in HeLa cells, whereas SMC3 knockouts resulted in a much more severe phenotype.

**Figure 5.**
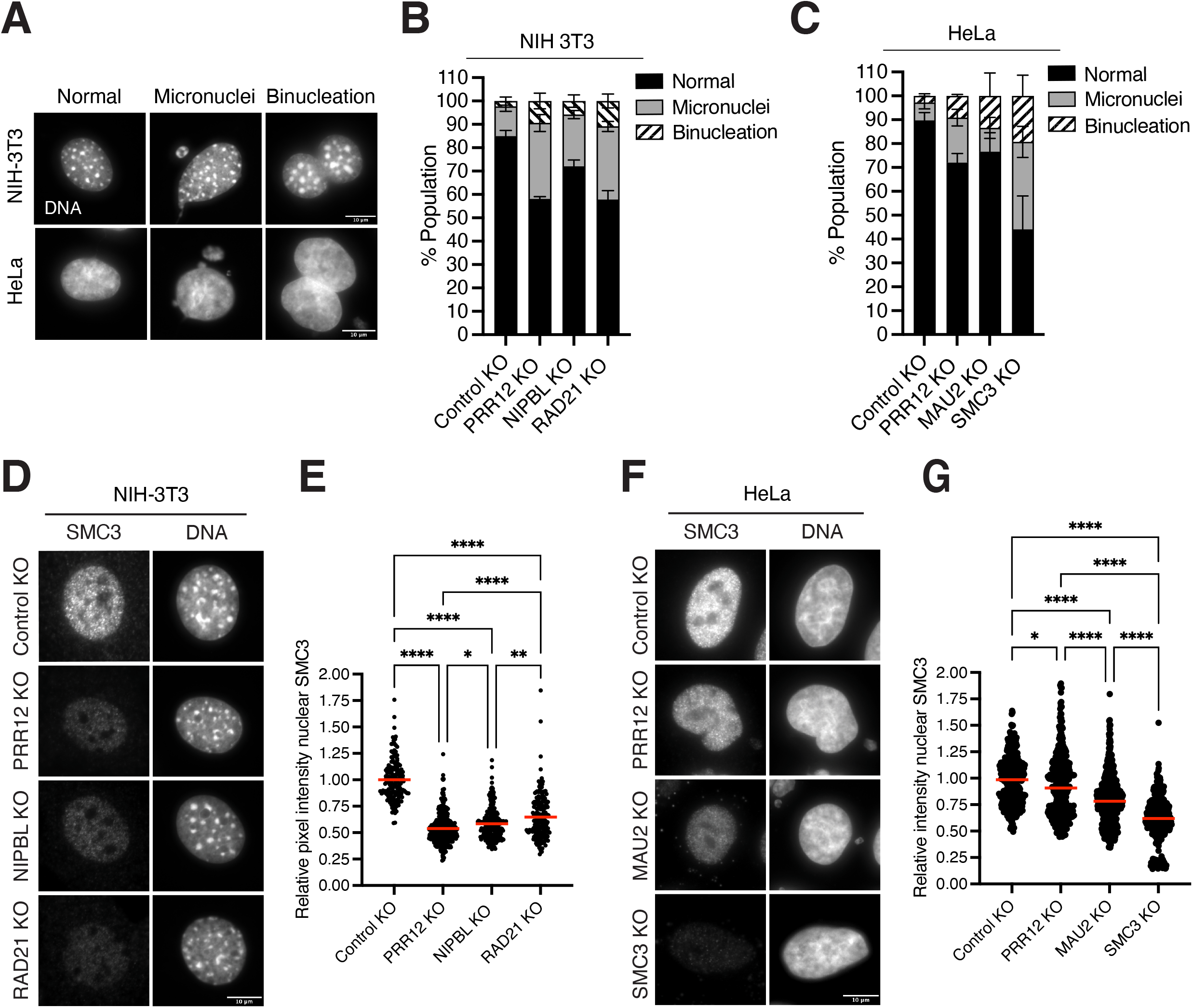
PRR12 is required for cohesin localization. (A) Representative Z-projected images of observed phenotypes (normal, macronuclei, binucleation) in mouse NIH-3T3 and human HeLa cell lines after knockout of the indicated genes (B-C). (B) Quantification of genome integrity phenotypes observed in the indicated NIH-3T3 cell lines, from A. *n* = approximately 500 cells per condition, across 3 experimental replicates. (C) Quantification of genome integrity phenotypes observed in the indicated HeLa cell lines, from A. *n* = approximately 500 cells per condition, across 3 experimental replicates. (D) Representative Z-projected images from knockout cells of the indicated genes in NIH-3T3 cell lines. Images show anti-SMC3 antibodies and DNA (Hoechst). (E) Quantification of relative pixel intensity nuclear SMC3 from D. Values are normalized to the average control value within each experiment. *n* = approximately 250 cells per condition, across 3 experimental replicates. One-way ANOVA was performed (* = 0.0306, ** = 0.0030, **** = <0.0001). (F) Representative Z-projected images from knockout cells of the indicated genes in HeLa cell lines. Images show anti-SMC3 antibodies and DNA (Hoechst). (G) Quantification of relative pixel intensity nuclear SMC3 from F. Values are normalized to the average control value within each experiment. *n* = approximately 300 cells per condition, across 3 experimental replicates. Error bars indicate SD. One-way ANOVA was performed (* = 0.0134,**** = <0.0001). Scale bars, 10 μM.

The loss of genome integrity is a core phenotype observed when cohesin is perturbed [62]. To determine if the loss of PRR12 impacts cohesin function, we analyzed cohesin localization using immunofluorescence. Using an antibody against SMC3, we found that the nuclear localization of cohesin was significantly decreased in NIH-3T3 PRR12 knockout cells (Fig. 5D-E), indicating PRR12 is required for the nuclear localization of cohesin. A similar reduction in nuclear-localized SMC3 was observed in NIPBL and RAD21 knockout NIH-3T3 cells (Fig. 5D-E). In contrast, PRR12 knockout in HeLa cells resulted in only a very modest decrease of nuclear localized SMC3 when compared to control knockout cells (Fig. 5F-G).

Our co-dependency predictions and protein interaction experiments indicated that PRR12 has a close connection with the NIPBL/MAU2 complex (Fig. 1B-C) [42, 60, 61]. NIPBL and MAU2 interact with a subset of cohesin complexes and are required for both cohesin DNA loop processivity and DNA entrapment [28, 31, 44–46]. NIPBL/MAU2 is capable of localizing to DNA independently of the cohesin complex (Fig. 5A-B) [36]. We therefore sought to test whether loss of PRR12 altered NIPBL localization. In mouse NIH-3T3 cells, knockout of PRR12 resulted in a modest decrease of NIPBL localization, suggesting NIPBL is sensitized to the loss of PRR12, but is not dependent on PRR12 for its localization (Fig. 6A-B). In contrast, in human HeLa cells, loss of PRR12 resulted in a significant increase in nuclear-localized NIPBL (Fig. 6C-D), indicating the relationship between these proteins may differ between mouse and human cell types.

**Figure 6.**
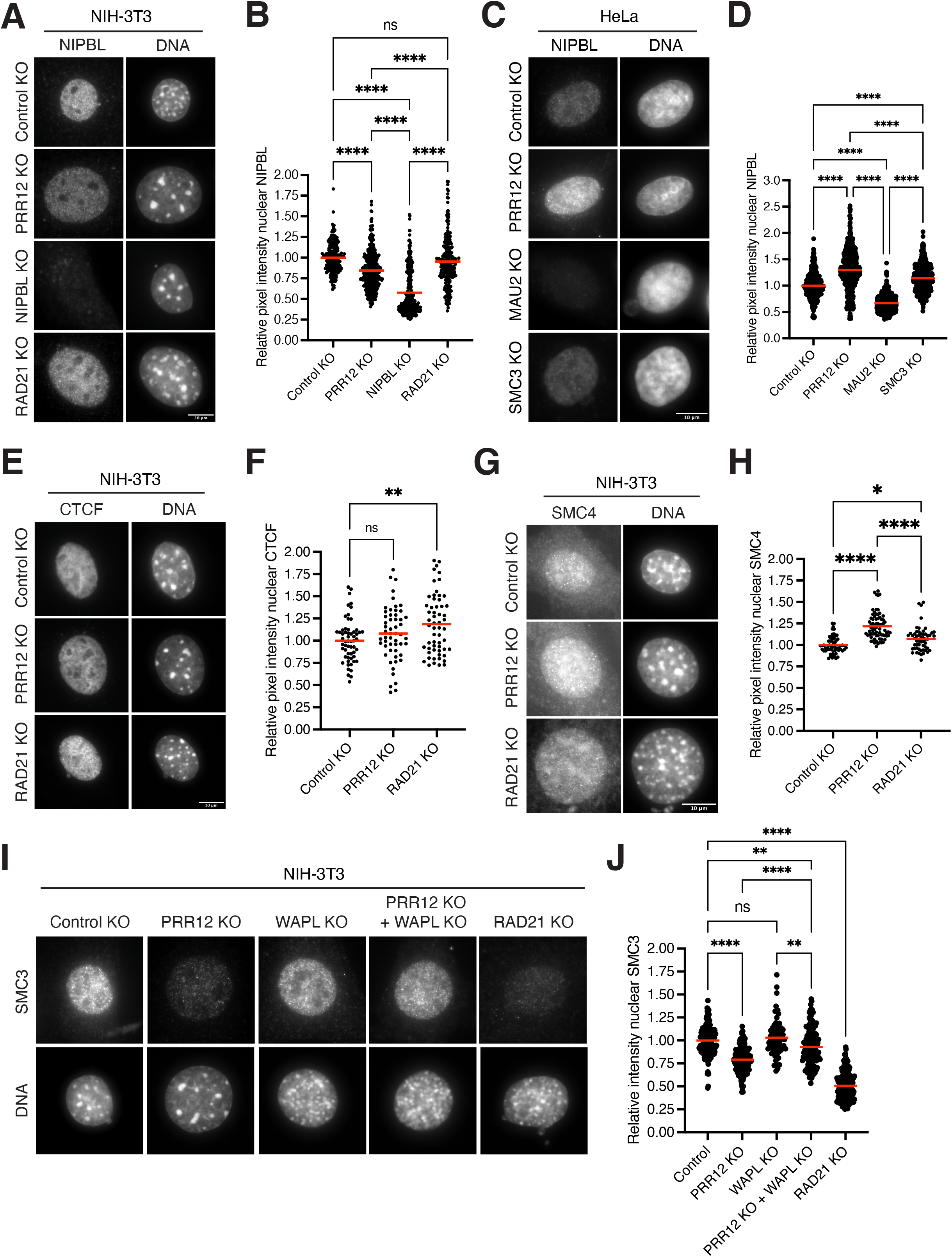
PRR12 contributes to cohesin stability. (A) Representative Z-projected images from knockout cells of the indicated genes in mouse NIH-3T3 cell lines. Images show anti-NIPBL antibodies and DNA (Hoechst). (B) Quantification of relative pixel intensity nuclear NIPBL from A. Values are normalized to the average control value within each experiment. *n* = approximately 250 cells per condition, across 4 experimental replicates. One-way ANOVA was performed (**** = <0.0001). (C) Representative Z-projected images from knockout cells of the indicated genes in human HeLa cell lines. Images show anti-NIPBL antibodies and DNA (Hoechst). (D) Quantification of relative pixel intensity nuclear NIPBL from C. Values are normalized to the average control value within each experiment. *n* = approximately 200 cells per condition, across 3 experimental replicates. One-way ANOVA was performed (**** = <0.0001). (E) Representative Z-projected images from knockout cells of the indicated genes in mouse NIH-3T3 cell lines. Images show anti-CTCF antibodies and DNA (Hoechst). (F) Quantification of relative pixel intensity nuclear CTCF from E. Values are normalized to the average control value within each experiment. *n* = approximately 50 cells per condition, across 2 experimental replicates. One-way ANOVA was performed (** = 0.0063). (G) Representative Z-projected images from knockout cells of the indicated genes in NIH-3T3 cell lines. Images show anti-SMC4 antibodies and DNA (Hoechst). (H) Quantification of relative pixel intensity nuclear SMC4 from G. Values are normalized to the average control value within each experiment. *n* = approximately 50 cells per condition, across 2 experimental replicates. One-way ANOVA was performed (* = 0.0421, **** = <0.0001). (H) Representative Z-projected images from knockout cells of the indicated genes in NIH-3T3 cell lines. Images show anti-SMC3 antibodies and Hoechst (DNA). (I) Quantification of relative pixel intensity nuclear SMC3 from H. *n* = approximately 200 cells per condition, across 3 experimental replicates. Error bars indicate SD. One-way ANOVA was performed (Control KO x PRR12 + WAPL KO ** =0.0086, WAPL KO x PRR12 + WAPL KO * =, 0.0035, **** = <0.0001). Scale bars, 10 μM.

Our data indicates PRR12 contributes to the nuclear localization of cohesin in mouse NIH-3T3 cells (Fig. 5D-E). To test whether this effect is specific to cohesin or could reflect a more global change in genome organization, we next sought to test the localization of other established genome organizers. We first quantified the localization of CTCF, a protein important for directing the localization of the cohesin complex, but which does not rely on cohesin for its nuclear localization (Fig. 6E-F) [14, 53, 63]. In contrast to the robust loss of cohesin in PRR12 knockout NIH-3T3 cells, CTCF levels were not significantly changed in PRR12 knockout cells compared to controls. Similarly, we assessed the localization of SMC4, a component of the condensin complex that mediates DNA compaction, and observed only very modest differences between control, PRR12, and RAD21 knockout NIH-3T3 cells (Fig. 6G-H).

The loss of cohesin in the absence of PRR12 suggests that PRR12 might be important for the loading or stabilization of cohesin on the DNA. To assess whether PRR12 contributes to cohesin stability, we tested the consequences of targeting WAPL, a cohesin destabilizer that acts to remove chromatin-bound cohesin. Prior work found that the co-depletion of WAPL and NIPBL restored the net levels of cohesin complexes on DNA [64]. In contrast to PRR12 knockout NIH-3T3 cells, in which nuclear localized SMC3 was significantly reduced, we found that simultaneous knockout of WAPL and PRR12 significantly increased the amount nuclear SMC3 to levels similar to control cells (Fig. 6I-J). These data suggest PRR12 may function to help stabilize DNA-bound cohesin complexes, counteracting WAPL-mediated cohesin removal. Together, these data implicate PRR12 as a novel regulator of cohesin.

### PRR12 is required for genomic integrity

In our functional analysis above, we found that PRR12 NIH-3T3 knockout cells appeared to have an increased nuclear size, consistent with altered genome behavior or a cell cycle arrest. To test cell cycle progression directly, we used FACS analysis of DNA content to define the proportion of cells in G0/G1, S phase, and G2/M. In contrast to control cells (G0/G1=43.5%, S =37.6%, G2/M =18.9%), we observed significantly more PRR12 knockout cells in G2/M (G0/G1 =48.4%, S=12.3%, G2/M=39.3%) with few S phase cells (Fig. 7A). Staining of cells with phospho-histone H3 (serine 10), a marker of mitosis, indicated that these cells were primarily in G2, with few mitotic cells in the absence of PRR12. SMC3 knockout cells also exhibited an increase in the proportion of G2/M cells (G0/G1=65.7%, S=4.2%, G2/M=30.1%) compared to controls, whereas NIPBL knockout cells contained a higher proportion of cells in G1 phase (G0/G1=63.5%, S=21.8%, G2=14.7%). In contrast, we did not observe a cell cycle arrest in PRR12 knockout or MAU2 knockout HeLa cells (Supplemental Fig. 2A), consistent with the differences observed in competitive growth assays in human and mouse cells for these genes (Fig. 4C-D). It is important to note that HeLa cells lack p53 function, which could account for the differences in cell cycle behaviors observed across the cell types when PRR12, NIPBL/MAU2, or cohesin are lost. However, p53 positive cells in the DepMap database also do not display a strong requirement for PRR12 function.

**Figure 7.**
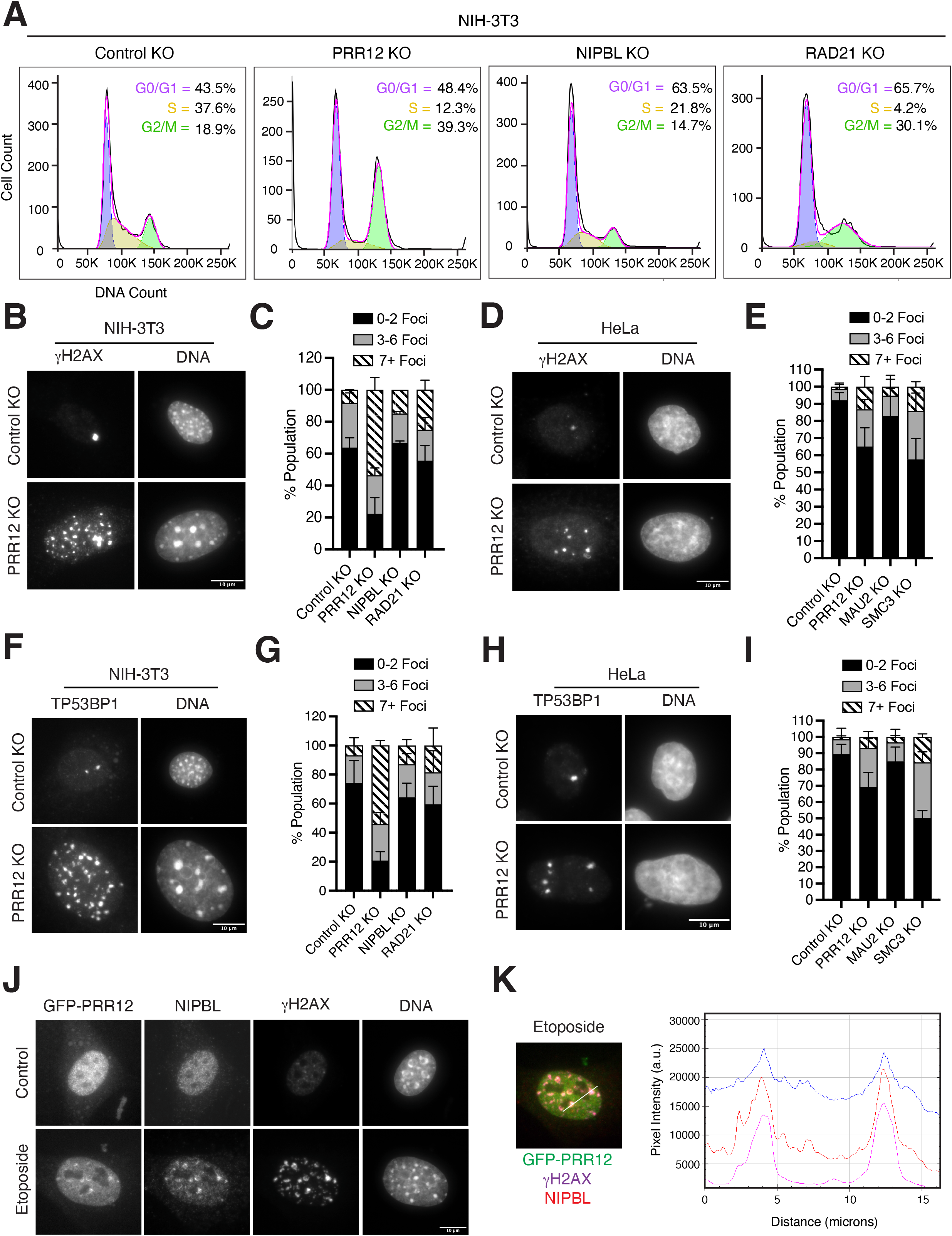
PRR12 is required for genomic integrity. (A) Representative DNA content analysis from the indicated knockout cell lines in mouse NIH-3T3 cells. Cells were fixed and stained for Hoechst and analyzed using flow cytometry. Cell stage was determined using Dean-Jett-Fox model, with equal gating set to all experimental groups. The percent population of cells in each cell cycle stage are indicated. *n* = approximately 10,000 cells analyzed per condition for each experiment, across 3 experimental replicates. (B) Representative Z-projected images from knockout cells of the indicated genes in NIH-3T3 cell lines. Images show anti-γH2AX antibodies and DNA (Hoechst). (C) Quantification of % population with foci from B. Fills denote the number of foci/cell. Values are normalized to the average control value within each experiment. *n* = approximately 300 cells per condition, across 3 experimental replicates. (D) Representative Z-projected images from knockout cells of the indicated genes in human HeLa cell lines. Images show anti-γH2AX antibodies and DNA (Hoechst). (E) Quantification of % population with foci from D. Fills denote the number of foci/cell. Values are normalized to the average control value within each experiment. *n* = approximately 400 cells per condition, across 3 experimental replicates. (F) Representative Z-projected images from knockout cells of the indicated genes in NIH-3T3 cell lines. Images show anti-TP53BP1 antibodies and DNA (Hoechst). (G) Quantification of % population with foci from F. Fills denote the number of foci/cell. Values are normalized to the average control value within each experiment. *n* = approximately 300 cells per condition, across 3 experimental replicates. (H) Representative Z-projected images from knockout cells of the indicated genes in HeLa cell lines. Images show anti-TP53BP1 antibodies and DNA (Hoechst). (I) Quantification of % population with foci from H. Fills denote the number of foci/cell. Values are normalized to the average control value within each experiment. *n* = approximately 250 cells per condition, across 2 experimental replicates. (J) Representative Z-projection of GFP-PRR12 expressing cells incubated in DMSO control or 1μM etoposide for 24 hours. Cells were fixed and stained for H2AX, NIPBL, and DNA (Hoechst). (K) Merged channels from the 1 μM etoposide treated cell from J, with line profile indicating pixel intensities of the denoted imaging channels. Error bars indicate SD.

The accumulation of cells in G2 in the absence of PRR12 suggests these cells may be undergoing a G2 checkpoint-mediated cell cycle arrest, which acts to prevent cells from entering mitosis in the presence of DNA damage or incomplete DNA replication [65–67]. We therefore sought to test whether PRR12 knockout cells exhibit DNA damage that could trigger the G2 checkpoint. To this end, we quantified the presence of DNA damage response factors in the absence of PRR12. We found that PRR12 knockout NIH-3T3 cells displayed a significant increase in the number of cells containing foci for multiple DNA damage response proteins, including γH2AX, TP53BP1, and RAD51B, compared to control cells (Fig. 7B-C, F-G, Supplemental Fig. 2B-C). Knockout of either NIPBL or RAD21 in NIH-3T3 cells also resulted in an increase in the accumulation of γH2AX, TP53BP1, and RAD51B foci, although this effect was milder compared to PRR12 knockout cells (Fig. 7B-C, F-G, Supplemental Fig. 2B-C). Similarly, knockout of PRR12, MAU2, and SMC3 resulted in an accumulation of DNA damage response proteins in HeLa cells, but to a lesser extent overall to that observed in NIH-3T3 cells (Fig. 7D-E, H-I). The presence of TP53BP1 and RAD51B foci are indicative of the presence of DNA double strand breaks in the absence of PRR12. Finally, to determine if the G2 checkpoint is activated in cells lacking PRR12, we analyzed phosphorylation of ATM at Ser1981 (pATM) (Supplemental Fig. 2D-E), which only becomes phosphorylated in response to DNA lesions [68, 69]. In PRR12 knockout NIH-3T3 cells, we observed the presence of pATM foci, consistent with the observed cell cycle arrest. Together, these data indicate that PRR12 is required for genomic integrity in both human and mouse cell types.

### PRR12 co-localizes with NIPBL at sites of DNA damage

In the absence of PRR12, cells accumulate spontaneous DNA double-strand breaks, eventually resulting in a G2 checkpoint-mediated arrest (Fig. 7A). Our mass spectrometry analysis revealed a strong interaction between PRR12 and multiple regulators of DNA damage across all tested samples (Fig. 3A), suggesting that PRR12 may function in the repair of DNA lesions. To further test this hypothesis, we assessed the localization of PRR12 in the presence of DNA damage. In control cells, GFP-PRR12 localized diffusely throughout the nucleus. In contrast, treatment with the DNA topoisomerase inhibitor etoposide resulted in an accumulation of GFP-PRR12 at DNA lesions, co-localizing with yH2AX (Fig. 7J-K). The cohesin complex is important for the regulation of DNA repair, specifically in the repair of double strand breaks by promoting homologous recombination [22, 24, 70, 71]. Previous work identified NIPBL as critical for the regulation of cohesin-mediated DNA damage repair, and NIPBL accumulates at DNA damage sites [72, 73]. Using an antibody specific to NIPBL, we found that NIPBL and GFP-PRR12 co-localized to sites of DNA damage marked by γH2AX, suggesting that PRR12 interacts with NIPBL at these sites. Together, these data indicate an essential role for PRR12 in maintaining genomic integrity and support a model in which PRR12 may function alongside NIPBL and cohesin in the repair of DNA lesions.

## Discussion

Here, we exploited the context-specific requirements inherent for cohesin functional diversity in combination with co-essentiality analysis to identify cohesin-regulatory proteins. Using this approach, we identified the previously uncharacterized gene PRR12 as a regulator of cohesin. In the absence of PRR12, cells display a significant reduction in nuclear-localized cohesin, a phenotype that can be rescued upon co-depletion of the cohesin destabilizer protein WAPL, suggesting PRR12 may contribute to the stability of cohesin on DNA. Furthermore, our affinity purifications revealed that PRR12 interacts with the cohesin complex via its C-terminal domain and we predict these interactions are mediated via an association with the NIPBL/MAU2 complex. Although similar phenotypes are observed in mouse NIH-3T3 cells upon NIPBL or PRR12 knockout, the severity of these phenotypes differs. These data suggest that the function of PRR12 does not overlap entirely with NIPBL. Additionally, although PRR12 displays strong co-dependencies with the NIPBL/MAU2 complex across cell lines, the requirements for these proteins do differ in some situations, indicating that NIPBL/MAU2 and PRR12 do not function as an obligate complex. Together, these data in combination with our affinity purification analysis, suggest that PRR12 interacts with multiple cohesin subunits and regulatory proteins that together contribute to shared and distinct cohesin functions.

In addition to a loss of nuclear localized cohesin, we found that PRR12 knockout cells accumulate DNA damage. Cohesin function is critical to the repair of DNA lesions [22, 25, 70, 71], and the localization of the cohesin complex to sites of DNA damage occurs in a NIPBL/MAU2-dependent manner [72, 73]. Our data support a model in which PRR12 may function together with NIPBL in mediating the response to DNA damage. In support of this model, we observed the co-localization of GFP-PRR12 with NIPBL at DNA lesions and our affinity purifications identified interactions between PRR12 and multiple DNA damage response proteins. PRR12 null cells have also been found to have an increased sensitivity to genotoxic agents in large-scale screens across multiple human cell lines [74, 75]. Reciprocally, the increased expression of PRR12 in large-scale CRISPR activation (CRISPRa) screens conferred resistance to DNA damage-inducing agents in multiple human cell lines [76, 77]. Alternatively, it is possible that the accumulation of spontaneous DNA breaks in the absence of PRR12 is a direct consequence of PRR12 loss. For example, PRR12 could contribute to DNA replication or function to relieve topological stress. Indeed, most spontaneous DNA double strand breaks occur during DNA replication [78] and our work identified interactions between PRR12 and the entire MCM replicative helicase complex. Cohesin also plays a role in replication fork progression and recent work has found that the cohesin-associated protein STAG2 is particularly important for this role [17, 18, 29, 79–82]. It is important to note that, similar to the dual role for cohesin in both DNA replication and the mediation of DNA damage, a role for PRR12 in both protecting against and response to DNA lesions either alongside cohesin or in addition to its cohesin associated roles does not need to be mutually exclusive.

Our work also demonstrates that the requirement for PRR12 differs across cell lines, with mouse NIH-3T3 cells significantly sensitized to PRR12 loss in comparison to human HeLa cells. Indeed, in most tested human cell lines, PRR12 is not considered essential for cell proliferation or survival based on large-scale CRISPR screens [42], with only select human cell lines displaying a growth defect in the absence of PRR12. However, in multiple mouse cell lines, loss of PRR12 results in significant growth defects (Fig.4D, Fig. 5A-B) [83]. Despite this significant difference in the requirement for PRR12, the protein sequence for PRR12 is highly conserved across species, with 93.6% identity between human and mouse cells. In addition, we observed similar localization dynamics and protein interactions for PRR12 in both human and mouse cell lines, suggesting the observed differential requirements do not reflect differences in PRR12 properties across species. Instead, these data most likely reflect differences in the underlying sensitivities of different cell lines to gene loss that may be influenced by the presence of functional redundancies, gene expression differences, altered sensitivities to stress, or overall differences in genome architecture. In fact, there a similar differential requirement for many cohesin regulatory proteins, with mouse BA/F3 and NIH-3T3 cells displaying increased sensitivity to gene loss compared to similar human cell types (Fig. 4B-D)[42].

Despite a non-essential role in cultured human cell lines, disruption of PRR12 function in human patients results in severe developmental and neurological impairments. Twenty-five distinct mutational variants in PRR12 have been identified across 24 patients, all of who present with developmental delays [47–50]. All of the documented variants are heterozygous mutations that occurred de novo. Predicted loss-of-function variants in PRR12 are extremely rare according to the Genome Aggregation Database (gnomAD(v3 and v2.1) suggesting that PRR12 is highly intolerant to haploinsufficiency [50]. Mutation in core cohesin regulators, including NIPBL, result in similar developmental and neurological defects, broadly defined as cohesinopathies. How disruption in NIPBL or PRR12 gene function manifest in disease pathologies remains poorly defined. The observed differences in gene requirements for PRR12 and known cohesin regulators in cell lines and human patients highlight the importance of studying the function of these genes in their native contexts. Together, our study highlights the power of identifying the cellular situations in which these genes are most required as a tool to reveal their function and to identify new players in these pathways. Harnessing these differences will provide a means to define the complex mechanisms by which cohesin is regulated as well as providing critical insight into how disruption of these processes drives disease.

## Acknowledgements

We thank Ana Losada for her support, insights, and guidance as well as members of the Cheeseman lab for their input and suggestions. This work was supported by grants from NSF (2029868) and NIH/NIGMS (R35GM126930) to IMC, a Damon Runyon post-doctoral fellowship to ALN, an American Cancer Society post-doctoral fellowship ES.

## Author Contributions

Conceptualization – ALN, IMC; Methodology – ALN, ES, IMC, Validation – ALN, ES, Investigation ALN, ES; Writing - Original Draft Preparation ALN, IMC; Writing – Review & Editing -ALN, ES, IMC; Visualization - ALN, ES; Supervision: IMC; Funding Acquisition: IMC.

## Materials and Methods

### Cell culture and Small Molecules

All HeLa and NIH-3T3 (ATCC-1568) cell lines were cultured in DMEM supplemented with 10% tetracycline-free fetal bovine serum (FBS), 100 units/ml penicillin, 100 units/ml streptomycin, and 2 mM l-glutamine (complete media) at 37°C with 5% CO_2_. Cell lines were tested monthly for mycoplasma contamination and validated based on the presence of appropriate cellular behaviors and markers. Polyclonal cell lines stably expressing N-terminal mEGFP fusions for human PRR12, or C-terminal PRR12 (1258-2036) under the expression of the EF1α promoter, were generated using lentiviral infection in HeLa and NIH-3T3 cells. GFP positive cells were enriched by FACS 2 days after transduction and cultured as described above. For cell stage synchronization, cells were incubated for 24 hours in the following small molecules: Palbociclib 1 μM (G1 phase; Selleckchem), Thymidine 2 mM (S phase; Sigma-Aldrich), RO-336 9 μM (G2 phase; Selleckchem). For analysis of GFP-PRR12 at DNA damage sites, cells were incubated in 1 μM Etoposide (MilliporeSigma) for 24 hours.

To ensure the robust, but conditional elimination of the target genes, we used a timed lentiviral infection strategy in which plasmids containing both Cas9 and the sgRNA targeting the gene of interest were transduced into cells. In brief, lentiviral plasmids containing pLCv2-opti-sgGeneX-mCherry, pLCv2-opti-sgGeneX-mTurquoise, were transduced into HeLa or NIH-3T3 cells by lentiviral transduction with spinfection (2250 RPM, 45 mins). Lentivirus was removed 16 hours after spinfection. At 2 days after transduction, successfully transduced cells were enriched by FACS (mCherry, mTurquoise) and cells were seeded for downstream experimental analysis. For knockout of PRR12 and WAPL simultaneously, a sgRNA targeting WAPL was first cloned into pLenti-sgNA under the U6 promoter and stably transduced into NIH-3T3 cells followed by puro selection. This cell line was then used as the parental cell line for the timed lentiviral infection strategy described above. It is important to note that due to the nature of CRISPR/CAS9 gene knockout approach used in this study, a proportion of the population of cells in each group will successfully repair the DNA or be heterozygous for gene knockout. This will result in variations in knockout efficiency across experiments.

### Plasmid Generation

Plasmids pLCv2-opti-stuffer-mCherry and pLCv2-opti-stuffer-mTurq2 were generated as previously described [84]. Individual sgRNAs designed using IDTdna.com and CRISPOR were then cloned into the pLCv2-Opti scaffold as previously described [85]. A control sgRNA with a single target site within the AAVS1 safe harbor site was used as a control. The following oligonucleotides (integrated DNA technologies) were used: sgAAVS1 (CACC GGGGCCACTAGGGACAGGAT), sgPRR12 Human (CACCGAGGGCTCCCCACCACCCAG), sgPRR12 Mouse (CACCGAAGTGGCTGAAGGAAGCAG), sgMAU2 Human (CACC GATGTGGCCCAGAAGCACGA), sgNIPBL Mouse (CACCGCTGTACCTCAAGCAGGTGC), sgRAD21 Mouse (CACCGGAGAGCATCATCTCACCAA), sgSMC3 Human (CACC GATGAAGAAAGGAGATGTGG). Codon optimized hPRR12 was synthesized at TWIST. Plasmids were assembled by polymerase chain reaction (PCR) amplification of inserts with Q5 DNA polymerase followed by digestion with restriction enzymes and ligation by T4-ligase into a Lentiviral backbone vector under the EF-1α promoter.

### Lentivirus Preparation and Transduction

For lentiviral production, HEK-293T cells were seeded at a density of 375,000 cells/mL in 2 mL complete media in 6-well plates. After 24 hours, cells were transfected with a mix containing 1.2 μg LentiVirus plasmid, 6 uL X-tremegene-9 transfection reagent (Roche 06365787001), 0.3 μg pCMV-VSV-G (Addgene Plasmid #8454), and 1 μg psPAX2 (Addgene plasmid #12260) in 50 μL Opt-MEM (Thermo Fisher Scientific). Media was changed 16 hours later to 1.5 mL fresh complete media. At 48 hours after transfection, virus was collected and stored at −80° C until use.

### Competition Assay

To assess relative cellular fitness, cells expressing pLCV2-opti-sgGeneX-mCherry were mixed 1:1 with cells expressing pLCV2-opti-sgAAVS1-mTurquose. Proliferation of the mCherry and mTuquoise-expressing cells was monitored by flow cytometry on an LSR Fortessa (BD Biosciences) flow cytometer. The percentage of mCherry-positive and mTurquoise-positive cells at the indicated timepoints was quantified and normalized to control levels at the same timepoint.

### FACS DNA content

For analysis of DNA content by flow cytometry, cells were fixed in 70% ethanol on ice for > 1 hour. Cells were washed 2 times in wash buffer (PBS + 0.1% Triton X100 + 3% BSA + 0.02% NaN3) followed by incubation for 1 hour in primary antibody in AbDil with rocking at 4°C. Cells were washed 3 times in wash buffer and incubated in CY5-conjugated antibodies at 1:300 and 0.6 mg/mL Hoescht-33342 (Invitrogen) for one hour with rocking at 4°C. Cells were washed 3 times in wash buffer, and resuspended in 500 uL AbDil for analysis by flow cytometry on the BD FACSCanto II (BD Biosciences) flow cytometer. Data were analyzed using FlowJo.

### Immunofluorescence microscopy

For immunofluorescence-based analysis, cells were plated on polylysine-coated coverslips (Sigma-Aldrich) and cultured for at least 24 hours prior to fixation. For timed lentiviral gene knockout experiments, cells were fixed approximately 3.5-4 days post viral transduction. Cells were fixed in 4% formaldehyde in PBS + 0.5% Triton X-100 for 10 mins followed by three consecutive washes in PBS + 0.1% Triton X-100 and blocked in AbDil (20 mM Tris-HCl pH 7.5, 150 mM NaCl, 0.1% Triton X-100, 3% BSA) for 30 min at room temperature. Immunostaining was performed by incubating the coverslips with primary antibodies diluted in AbDil for 1 hour at room temperature followed by three consecutive washes in PBS + 0.1% Triton X-100. After washing, secondary antibodies were diluted 1:300 in Abdil together with 0.3 μg/mL Hoescht-33342 (Invitrogen) and the sample was incubated for 1 h at room temperature followed by 3 consecutive washes in PBS containing 0.1% Triton X-100. The coverslips were next mounted in p-phenylamine diamine (PPDM) onto coverslips. Immunofluorescence cell images were acquired on a DeltaVision Core deconvolution microscope (Applied Precision) equipped with a CoolSnap HQ2 CCD camera. Approximately 35 Z-sections were acquired at 0.2 μm steps using a 60×, 1.42 NA Olympus U-PlanApo objective.

The following antibodies were used: Phospho-ATM (Ser1981) (10H11.E12) (1:500, Cell Signaling; 4526), PRR12 (1:250, Abcam; ab121354), SMC3 (C.terminal) (1:500, Abcam; ab155587), NIPBL (1:500, gift from Ana Losada), Recombinant Gamma H2A.X (phosphor S139 [n1-431] (1:500, Abcam; ab303656), RAD51B (1:500, Thermo Fischer; MA525683), tP53BP1 (1:500, VWR; 102108-436), CTCF (1:500, Cell Signaling; D31H2), SMC4 (1:500, Cell Signaling; D14E2), H3S10p (1:250; Abcam: ab5176).

### Immunoprecipitation–mass spectrometry

HeLa or NIH3T3 cells that stably express either a mEGFP-hPRR12 or a mEGFP-hPRR12 (1258-2036) were cultured as described above. Cells were harvested by incubation in PBS supplemented with 5 mM EDTA for 5 minutes at 37°C. Harvested cells were pelleted by centrifugation at 200 x g. The cell pellets were then washed once with 1X PBS and once with 1x lysis buffer (25 mM HEPES pH 8.0, 2 mM MgCl_2_, 0.1 mM EDTA pH 8.0, 0.5 mM EGTA pH 8.0, 300 mM KCl and 10% glycerol). Cells were pelleted by centrifugation at 200 x g after each wash step. Next, the cells were resuspended at a 1:1 ratio in 1x lysis buffer and drop frozen in liquid nitrogen before being stored at −80°C. 1.5x lysis buffer (37.5 mM HEPES pH 8.0, 3 mM MgCl2, 0.15 mM EDTA pH 8.0, 0.75 mM EGTA pH 8.0, 450 mM KCl, 15% glycerol, and 0.075% NP-40) supplemented with 1 Complete EDTA-free protease inhibitor cocktail (Roche), 20 mM beta-glycerophosphate, 1 mM sodium fluoride, and 0.4 mM sodium orthovanadate was added. After addition of 1.5x lysis buffer cells were thawed in a 37°C water bath and 1 mM PMSF was added. Cells were then lysed by sonication and the lysate was clarified by centrifugation. Clarified lysates were incubated on a rotating wheel with NHS Mag Sepharose beads (Cytiva) coupled to a GFP nanobody at 4°C for 1 hour. After incubation with the clarified lysate the beads were rinsed 3 times with 1x lysis buffer supplemented with 0.05% NP-40, 1 mM DTT, and 10 μg/ml leupeptin/pepstatin/chymostatin and then washed 2 times for 5 minutes on a rotating wheel at 4°C in the same buffer. Then cells were washed once in 1x lysis buffer supplemented with 1 mM DTT, and 10 μg/ml leupeptin/pepstatin/chymostatin. Next, bound proteins were eluted by washing the beads with 2 bead volumes of 100 mM glycine pH 2.6 for 5 minutes on a rotating wheel. Three separate elutions were pooled and Tris pH 8.5 was added to a final concentration of 200 mM. The eluted proteins were precipitated at 4°C overnight by the addition of 1/5 the final volume of trichloroacetic acid. The precipitated proteins were pelleted by centrifugation and washed 2 times with cold acetone. Next, the pelleted proteins were resuspended in 5% SDS, 50 mM tetraethylammonium bromide pH 8.5 (TEAB), 20 mM DTT before being incubated at 95 °C for 10 minutes. The proteins were allowed to cool to room temperature before the addition of 40 mM iodoacetamide. The proteins were incubated with iodoacetamide for 30 minutes in the dark and then 2.5% v/v phosphoric acid was added. Next, 6 times the volume of S-trap binding buffer (90% MeOH, 100 mM TEAB, pH 7.55) was added to the acidified proteins before the proteins were loaded onto a S-Trap microcolumns (ProtiFi). The column was washed 4 times with S-trap binding buffer and then 1 μg of trypsin diluted in 50 mM TEAB pH 8.5 was added to the column and the proteins were digested on-column overnight at 37°C in a humidified incubator. The digested peptides were then eluted with three separate elutions: 50 mM TEAB, 0.2% formic acid then 50% acetonitrile/0.2% formic acid. The elutions were pooled and quantified with a fluorometric peptide assay kit (Pierce) and then the eluted peptides were lyophilized overnight. Lyophilized peptides were resuspended in 0.1% formic acid and then injected onto a Orbitrap Eclipse Tribrid (Thermo Fischer Scientific) connected to a Vanquish Neo UHPLC system. Proteins were identified in Proteome Discoverer 2.4 (Thermo Fisher Scientific) using Sequest HT. Percolator was used to validate Peptide-spectrum matches.

### Quantification and Statistical Analysis

Quantification of fluorescence intensity was conducted on unprocessed, maximally projected images using FIJI/image J and Cell Profiler. For image quantification, all images for comparison were acquired using the same microscope and acquisition settings. Statistical analyses were performed using Prism (GraphPad Software). Details of statistical tests and sample sizes are provided in the figure legends.

