## Supplemental Legends and Figures for "Co-Essentiality Analysis Identifies PRR12 as a Regulator of Cohesin and Genome Integrity"

**Supplemental Information Titles and Legends**

**Supplementary Figure 1: Related to Figure 2**
(A) Alphafold structural predictions of hPRR12 predicted binding domains. (B) Amino acid sequence of hPRR12 predicted binding domain 1. (C) Amino acid sequence of hPRR12 predicted binding domain 2.

**Supplemental Figure 2: Related to Figure 5**
(A) Representative DNA content analysis from the indicated knockout cell lines in human HeLa cells. Cells were fixed and stained for Hoechst and analyzed using flow cytometry. Cell stage was determined using Dean-Jett-Fox model, with equal gating set to all experimental groups. The percent population of cells in each cell cycle stage are indicated. *n* = approximately 10,000 cells analyzed per condition for each experiment, across 2 experimental replicates. B) Representative Z-projected images from knockout cells of the indicated genes in Mouse NIH-3T3 cell lines. Images show anti-RAD21 antibodies and DNA (Hoechst). (C) Quantification of % population with foci from B. Fills denote the number of Foci/cell. Values are normalized to the average control value within each experiment. *n* = approximately 250 cells per condition, across 3 experimental replicates. D) Representative Z-projected images from knockout cells of the indicated genes in Mouse NIH-3T3 cell lines. Images show anti-pATM (Ser1981) antibodies and DNA (Hoechst). (E) Quantification of % population with foci from D. Fills denote the number of Foci/cell. Values are normalized to the average control value within each experiment. *n* = approximately 250 cells per condition, across 3 experimental replicates.

**Supplementary Table 1**
Raw Data: GFP-hPRR12 affinity purification mass spectrometry results for GFP-PRR12 in Human Hela cells

**Supplementary Table 2**
Raw Data: GFP-C.Terminus hPRR12 affinity purification mass spectrometry results for GFP-PRR12 in Human Hela cells

**Supplementary Table 3**
Raw Data: GFP-hPRR12 affinity purification mass spectrometry results for GFP-PRR12 in Mouse NIH-3T3 cells

**Supplementary Table 4**
Raw Data: GFP-C.Terminus hPRR12 affinity purification mass spectrometry results for GFP-PRR12 in Mouse NIH-3T3 cells

**A**

**Alphafold prediction PRR12 (1551- 2036)**

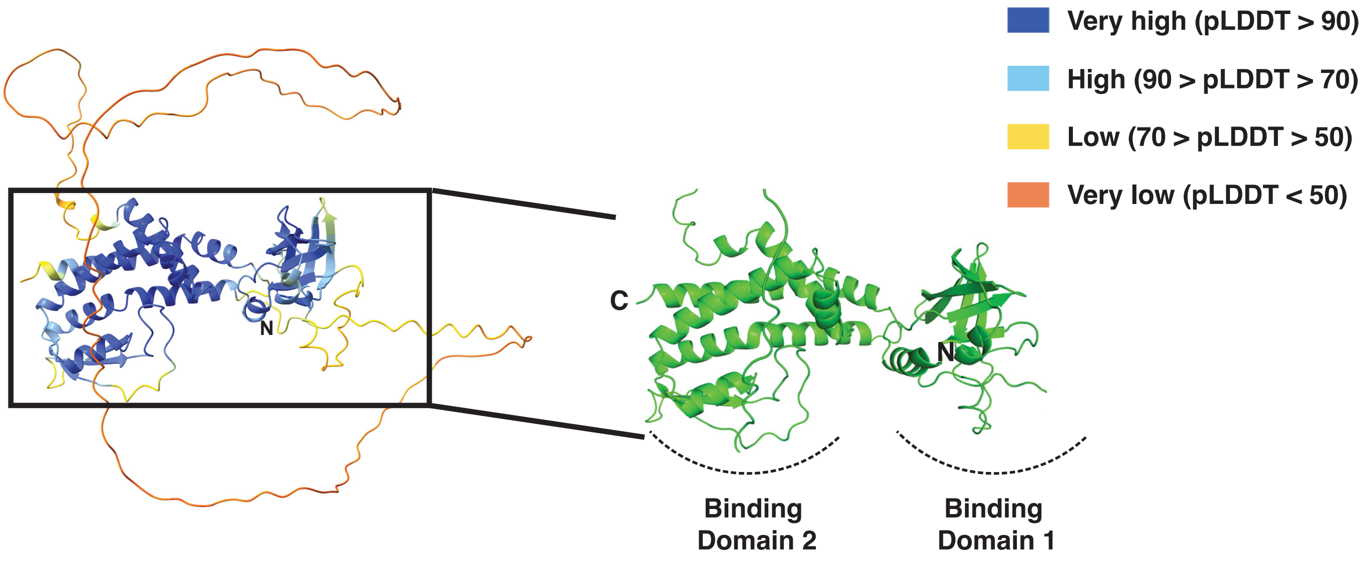


**B**

```
                   1570      1580      1590      1600      1610      1620
                    |         |         |         |         |         |
H. Sapiens    EAGESGGEGIFRERDEFVIRAEDIPSLKLALQTGREPPPIWRVQKALLQKFTPEIKDGQR     1623
M. Musculus   EAGESGGEGIFRERDEFVIRAEDIPSLKLALQTGREPPPIWRVQKALLQKFTPEIKDGQR     1617
X. Laevis     EDVESSGEGIFRERDEFVIRVEDIQALKLALQTGREPPPIWRVQKALLQKFTPEIKDGQR     1570
D. Rerio      DESESGGEGIFRERDEFVVRIEDIRSLKMALRTGREPPPIWRVQKALLQKFSPEIKDGQR     2124
              :  **.************:* *** :**:**:*******************:********

                   1630      1640      1650      1660      1670      1680
                    |         |         |         |         |         |
H. Sapiens    QFCATSNYLGYFGDAKNRYQRLYVKFLENVNKKDYVRVCARKPWHRPPVPVRRSGQAKNP     1683
M. Musculus   QFCATSNYLGYFGDAKNRYQRLYVKFLENVNKKDYVRVCARKPWHRPPLPVRRSGQTKGP     1677
X. Laevis     QFCATSNYLGYFGDAKNRYQRLYVKFLENVNKKDYVRVCSKKPWHRPLQTMRRQSQTKAP     1630
D. Rerio      QFCATSNYLGYFGDAKKRYQRLYVKFLENINKKDYVRVCSRKPWHRPSLNLRRQSQPVPK     2184
              ****************:************:*********::******   :**..*
```

**C**

```
             1840      1850      1860      1870      1880      1890
              |         |         |         |         |         |
H. Sapiens    LLKTRAMREMYRSYVEMLVSTALDPDMIQALEDTHDELYLPPMRKIDGLLNEHKKKVLKR     1899
M. Musculus   LLKTRAMREMYRSYVEMLVSTALDPDMIQALEDTHDELYLPPMRKIDGLLNEHKKKVLKR     1898
X. Laevis     VLNTRAMKEMFRSYIEMLVSTALDPDMIQALEDTSDELYLPPMRKIDGIVNEHKKKVLKK     2218
D. Rerio      GLNTRAMREMYRSYVEMLVSTALDPDMIQALEDTEDELYLPPMRKIDSIISEQKRKLLKR     2404
               *:****:**:***:******************* ************.::.*:*:*:**:

             1900      1910      1920      1930      1940      1950
              |         |         |         |         |         |
H. Sapiens    LSLSPALQDALHTFPQLQVEQSGEGSPEEGAVRLRPAGEPYNRKTLSKLKRSVVRAQEFK     1959
M. Musculus   LSLSPALQDALHTFPQLQVEQTGEGSPEEGAVRLRPAGEPYNRKTLSKLKRSVVRAQEFK     1958
X. Laevis     ISLSSSFQEAIHTFPQLNSE------PGEPSTRMKPGGEAYNRKTLNKLKKNVAKPQEFK     2272
D. Rerio      VNMNSQHQETLHTFPQITAEPL-----DSGLVRVRLGGECYNRKTLNRIKKSVPKPQDLK     2459
              :.:.   *:::*****:  *        .  .*:: .** ******.::*:.* : *::*

             1960      1970      1980      1990      2000      2010
              |         |         |         |         |         |
H. Sapiens    VELEKSGYYTLYHSLHHYKYHTFLRCRDQTLAIEGGAEDLGQEEVVQQCMRNQPWLEQLF     2019
M. Musculus   VELEKSGYYTLYHSLHHYKYHTFLRCRDQTLAIEGGAEDLGQEEVVQQCMRNQPWLEQLF     2018
X. Laevis     VDAEKSLFYTLYHSLHHYKYHTFLRCKQETNAIEEQNDDLGQEEVVQQCMRNQPWLEKLF     2332
D. Rerio      LSTEMCRIYSLYHSLHHYKYHTFLHCKKETNTIEQASEDPGQEEVVQQCMANQNWLETLF     2519
              :. * .  *:**************:*:.:* :**   :* ********** ** *** **

             2020      2030
              |         |
H. Sapiens    DSFSDLLAQAQAHSRCG          2036
M. Musculus   DSFSDLLAQAQAHSRCG          2035
X. Laevis     DSFIDLITQAQNKC--A          2347
D. Rerio      NSFLELMALSTKV----          2532
              :** :*:: :
```

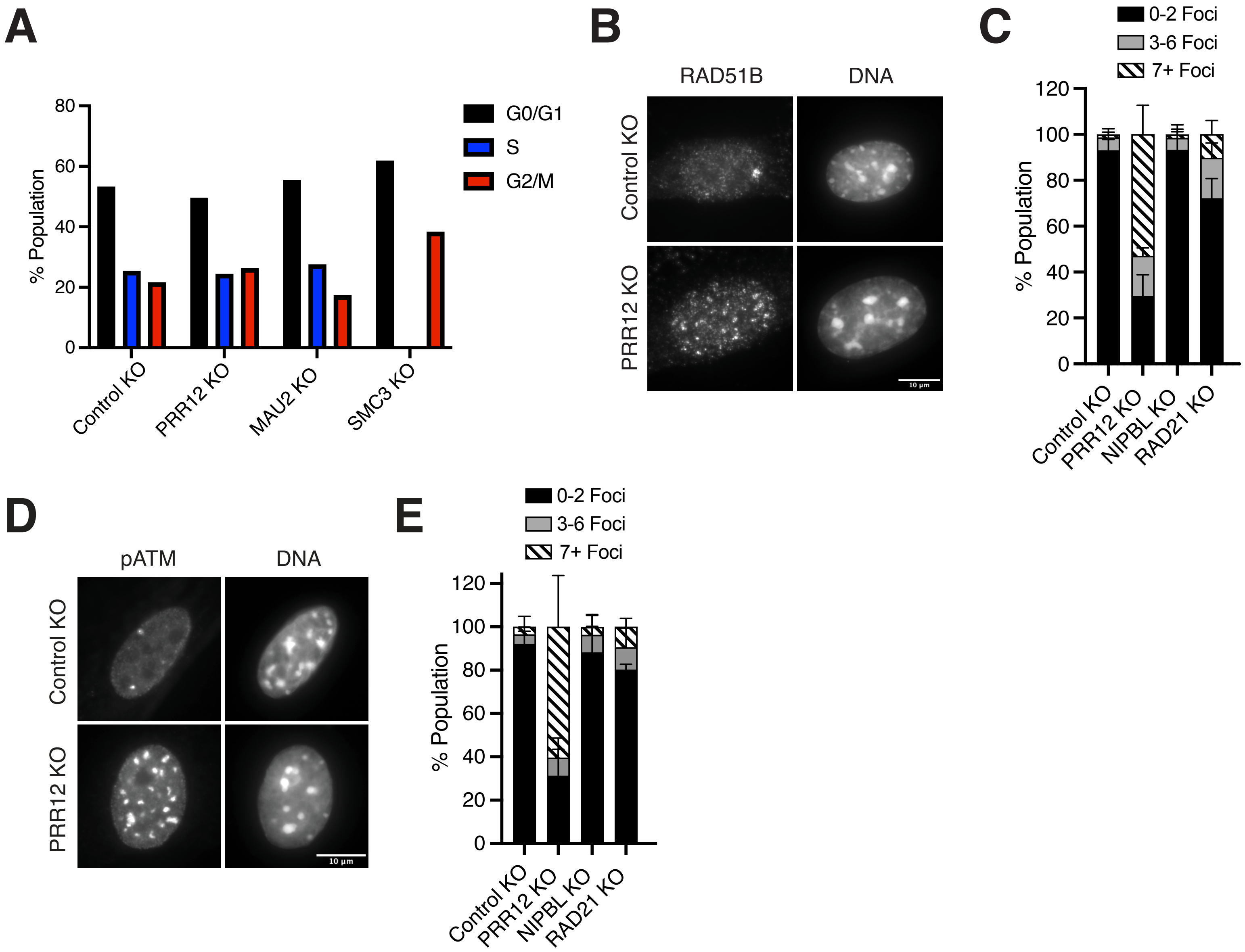
A
G0/G1
S
G2/M
% Population
80
60
40
20
0
Control KO
PRR12 KO
MAU2 KO
SMC3 KO
B
RAD51B
DNA
Control KO
PRR12 KO
10 µm
C
0-2 Foci
3-6 Foci
7+ Foci
% Population
120
100
80
60
40
20
0
Control KO
PRR12 KO
NIPBL KO
RAD21 KO
D
pATM
DNA
Control KO
PRR12 KO
10 µm
E
0-2 Foci
3-6 Foci
7+ Foci
% Population
120
100
80
60
40
20
0
Control KO
PRR12 KO
NIPBL KO
RAD21 KO
